# Comparative cytogenetics of kenaf (*Hibiscus cannabinus* L.) breeding lines reveal chromosomal variability and instability

**DOI:** 10.1101/2023.11.18.567672

**Authors:** Nii-Ayi Ankrah, Abdullah El-nagish, Sarah Breitenbach, Antonia Tetteh, Tony Heitkam

**Author notes:** Antonia Tetteh and Tony Heitkam share corresponding authorship.

## Abstract

Kenaf (*Hibiscus cannabinus*), a native warm-seasonal crop in Africa, is being considered for genetic improvement for local bast fiber production. To expedite its genetic improvement through breeding, kenaf genotypes from Ghana were assessed for genomic diversity regarding their chromosomal composition and ploidy levels. To gain insight into the repetitive DNA fractions in kenaf, the organization of 5S and 35S rRNA genes, as well as telomeric signal patterns were studied by a molecular cytogenetic approach.

Using multi-color fluorescent *in situ* hybridization, distinct rDNA loci and *Arabidopsis*-type telomeres were revealed. The 5S rRNA genes were conserved in kenaf and localized in interstitial regions of two chromosomes across all accessions. The 35S rRNA genes were variable across the kenaf accessions and localized at sub-terminal ends and rarely interstitially in eight or six chromosome arms. Telomeric signals were observed at terminal ends of all chromosomes, with smaller signals also interstitially. The chromosome configuration of Ghana kenaf accessions was confirmed to be 2n=2x=36, each. We discuss the chromosomal variability and the likely genomic instability in the kenaf breeding lines from Ghana.

To our knowledge, this is the first report on molecular cytogenetics on kenaf and thus, provides valuable insights into the genome of kenaf that will be useful for breeding. Additionally, this study provides a basis for further studies to analyze the repetitive DNA sequences and develop reference karyotypes to reveal genetic and evolutionary relationships between cultivated and wild *Hibiscus* species.

## INTRODUCTION

The *Hibiscus* genus of the family *Malvaceae* contains about 300 species of herbs, shrubs and trees that grow around the globe in tropical, sub-tropical and temperate regions (Jo et al. 2019). Kenaf (*Hibiscus cannabinus*) is a day-length sensitive dicotyledonous plant characterized by rapid growth that could reach a height of about 5 meters under favorable climatic conditions (Deng et al. 2017). Kenaf is considered an economic plant in Africa because of the bast fibers obtained from the stem, which has many applications including usage as ropes and cordages, fish nets, composites in building materials and electrical appliances, and as a jute substitute for making sacks for packaging of agricultural produces (Humphries 2004; Schmiedel et al. 2014; Muniandi et al. 2018; Zhang et al. 2020).

Due to the economic value of kenaf, making genetic improvements to the crop for breeding purposes is very important. Genetically, a kenaf reference genome has been sequenced (Zhang et al. 2020) and its major genomic hallmarks are known (genome size 1C = 1 Gb; chromosome configuration 2n=2x=36). Additionally, genetic linkage maps with varying linkage groups (Chen et al. 2008; Chen et al. 2011; Zhang et al. 2011; Wu et al. 2016; Zhang et al. 2020) have been constructed in kenaf. Hence, regarding the assessment of kenaf’s molecular diversity, most studies to date rely on molecular markers. These encompass simple and inter-simple sequence repeats (SSR, ISSR; Tao et al. 2005; Satya et al. 2013), randomly amplified polymorphic DNA (RAPD; Cheng et al. 2002; Faruq et al. 2015), amplified fragment length polymorphism (AFLP; Cheng et al. 2004; Coetzee et al. 2008; Kim et al. 2010), sequence-related amplified polymorphism (SRAP; Tao et al. 2005; Qi et al. 2011; Zhang et al. 2011), and single nucleotide polymorphism (SNP) markers (Zhang et al. 2020). Nevertheless, kenaf breeding is still impaired by the lack of genetic understanding of the kenaf germplasm and native breeding resources. Only the most basic facts are known about species of the genus *Hibiscus*, such as the occurrence of high ploidy levels up to hexadecaploidy (Menzel & Wilson 1969; Wilson 2006; Zhang et al. 2020; Luo & He 2021).

To better inform local breeding policies, the genomic diversity of kenaf and the kenaf accessions from the wild needs to be determined, especially regarding their chromosomal composition and ploidy levels. However, cytogenetics of kenaf genotypes was barely performed, and the current knowledge has been mostly obtained by fluorochrome staining (Hiron et al. 2006; Islam & Alam 2011) and karyotype analysis (Akpan & Hossain 1998). We argue that chromosome analysis by comparative fluorescence *in situ* hybridization (FISH) – a gold standard for molecular cytogenetics (Fishman et al. 2013; Murat et al. 2015; Heitkam & Garcia 2023) – would provide information on the chromosome characteristics and genomic variability of the species. For this, the ribosomal RNA genes constitute ideal cytogenetic markers as they are highly conserved and usually occupy large chromosomal territories along plant chromosomes (Ksiazczyk et al. 2010; Kolano et al. 2015).

Here, we describe an efficient protocol for chromosome preparation of kenaf from primary roots. To understand the chromosomal composition of kenaf breeding accessions, we perform FISH with tandemly repeated probes binding to the 35S and 5S ribosomal genes as well as the telomeres. We follow the number and localization of these signals across several breeding accessions to understand the chromosomal stability of kenaf breeding accessions and to provide a solid basis for further cytogenetic exploration of the kenaf wild germplasm.

## MATERIALS AND METHODS

### Plant material and genomic DNA extraction

Seeds of kenaf breeding lines and landrace were provided by the Bast Fiber Research Unit (KNUST, Ghana; see **Table 1**). These lines represent the typical variation in kenaf leaf morphology, including unlobed, tri-, penta- and hepta-lobed leaves (see **Table 1**; **Supplementary Figure 1**). Seedlings were grown under greenhouse conditions at the Faculty of Biology, TU Dresden, Germany. Genomic DNA was isolated from young leaves using the Machery-Nagel Genomic DNA Isolation Kit (Germany), according to the manufacturer’s protocol, however, genomic DNA was eluted with double distilled water (ddH_2_O) instead of the elution buffer provided with the isolation kit.

**Table 1:** Chromosome numbers and number of 5S and 35S rDNA sites of five kenaf accessions.

| Species | Accession number | Chromosome configuration | Number of rDNA sites |  |
| --- | --- | --- | --- | --- |
|  |  |  | 5S | 35S |
| *H. cannabinus* 21 (Land race) | HC-21 | 2n=2x=36 | 2 | 8 |
| *H. cannabinus* (Entire narrow leaves) | EN-31 | 2n=2x=36 | 2 | 8 |
| *H. cannabinus* (Trilobed narrow leaves) | TN-11 | 2n=2x=36 | 2 | 8 |
| *H. cannabinus* (Pentalobed narrow leaves) | PN-11 | 2n=2x=36 | 2 | 8 |
| *H. cannabinus* (Heptalobed narrow leaves) | HN-11 | 2n=2x=36 | 2 | 6 |

### Chromosome preparation

Kenaf seeds were germinated in petri dishes in the dark and after three days, primary roots were collected and incubated separately in 2 mM 8-hydroxyquinoline for 1.5 h and in nitrous oxide at 10.5 bar for 35 minutes at room temperature. The roots were immediately fixed in fixative solution (methanol 3:1 acetic acid) for 30 minutes, discarded, and fixed in fresh fixative and stored at 4 °C until further use.

Fixed roots were washed once in distilled water and twice in citrate (enzyme) buffer (4 mM citric acid– 6 mM sodium citrate, pH 4.6) for 5 minutes each. Roots were incubated at 37°C for 135 minutes in an enzyme mixture containing 5% (w/v) pectinase from *Aspergillus niger* (Sigma), 2% (w/v) cellulase Onozuka R 10 (Serva), 0.5% (w/v) pectolyase from *Aspergillus japonicus* (Fluka), and 2% (w/v) cytohelicase from *Helix pomatia* (Sigma) in citrate (enzyme) buffer (4 mM citric acid– 6 mM sodium citrate, pH 4.6). The digested root tips were macerated on pre-cleaned slides in a drop of citrate (enzyme) buffer using a dissecting needle. About 150 µl of 45% acetic acid was added and the mixture was incubated for 5 minutes at room temperature. Freshly prepared fixative solution (methanol 3:1 acetic acid) was added in drops to spread and fix the chromosomes on the slides. Slides were air-dried and examined for good mitotic chromosomes using a phase contrast microscope (Axioskop 40, Carl Zeiss).

### Labeling of probes, fluorescent *in situ* hybridization (FISH) and image acquisition

The physical mapping of rDNA sites involved the use of published probes 18S-1 and pXV1. The 18S-1 probe contains a 552-bp fragment of the sugar beet 18S-5.8S-25S ribosomal RNA (rRNA) gene with the 35S rDNA (Schmidt et al. 1994) labeled with digoxigenin-11-dUTP by PCR (Roche) and the probe pXV1 for the 5S rRNA gene (Schmidt et al. 1994) labeled with biotin-11-dUTP by PCR (Roche). The telomeric probe pLT11 from Arabidopsis (Richards & Ausubel 1988) was labeled with DY-415-aadUTP by PCR (Dyomics). PCR labeling was performed with standard M13 primers using the following program: 5 min 95°C, 35 cycles of (1 min 95°C, 40 sec 56°C, 1 min 72°C), 10 min 72°C.

Fluorescent *in situ* hybridization was performed according to Liedtke et al. (2022). Briefly, chromosome slides were pre-treated with 100 µg/mL RNase A in 2× SSC for 30 min at 37 °C and washed twice in 2× SSC. After incubation with 10 µg/mL pepsin in 0.01 mM HCl for 20 min at 37 °C, preparations were stabilized in freshly de-polymerized 4%(w/v) paraformaldehyde in double distilled water (ddH_2_O) for 15 min, washed again with 2× SSC, and dehydrated in an ethanol series (70%; 90%; 100%) followed by air-drying at room temperature. A 30 µL hybridization mixture containing 100% formamide, 20× SSC, 10% sodium dodecyl sulfate (SDS), 1 µg/µL salmon DNA (blocking DNA), 50% dextran sulfate, labeled FISH probes, and ddH_2_O was prepared per each metaphase slide. The hybridization mixture (30 µL) was denatured for 10 min at 70 °C and applied per slide. After the application of the hybridization mixture, the slides were covered with plastic coverslips, denatured at 70 °C for 8 min, cooled down stepwise to 37 °C in an *in situ* Omni-slide thermal cycler (Thermo Electron) and hybridized overnight at 37 °C in a humid chamber. Stringent post-hybridization washing was performed at 42 °C in 20 % formamide diluted with 4× SSC/0.2 % Tween 20. Biotin-labeled probes were detected by streptavidin coupled to Cy3. Digoxigenin-labeled probes were detected with anti-digoxigenin coupled to fluorescein isothiocyanate (FITC; Roche). Chromosome preparations were counter-stained with 4′,6-Diamidino-2-phenylindole (DAPI) in an antifade solution (Citifluor). Slides were examined with a fluorescence microscope (Zeiss Axioplan 2 Imaging) equipped with ultraviolet excitation filters; filter 01 (DAPI), filter 09 (FITC), and filter 15 (Cy3).

Images were acquired directly with the Applied Spectral Imaging v. 3.3 software coupled with the high-resolution CCD camera (ASI BV300-20A) and processed using Adobe Photoshop v. 7.0 using functions that affect the whole image equally after contrast optimization.

## RESULTS

### *H. cannabinus* breeding lines are diploid with 36 chromosomes

For obtaining good metaphases of kenaf, chromosome preparations were optimized using 8-hydroxyquinoline and nitrous oxide in a pressure chamber as separate treatments (Figure 1A, 1B). The results showed that kenaf roots treated with 2 mM 8-hydroxyquinoline for 80 min resulted in well-spread metaphases without chromatid separation (Figure 1A). For root treatment with nitrous oxide, the results showed well-spread metaphases with distinct separation of chromatids after 35 minutes (Figure 1B).

**Figure 1:**
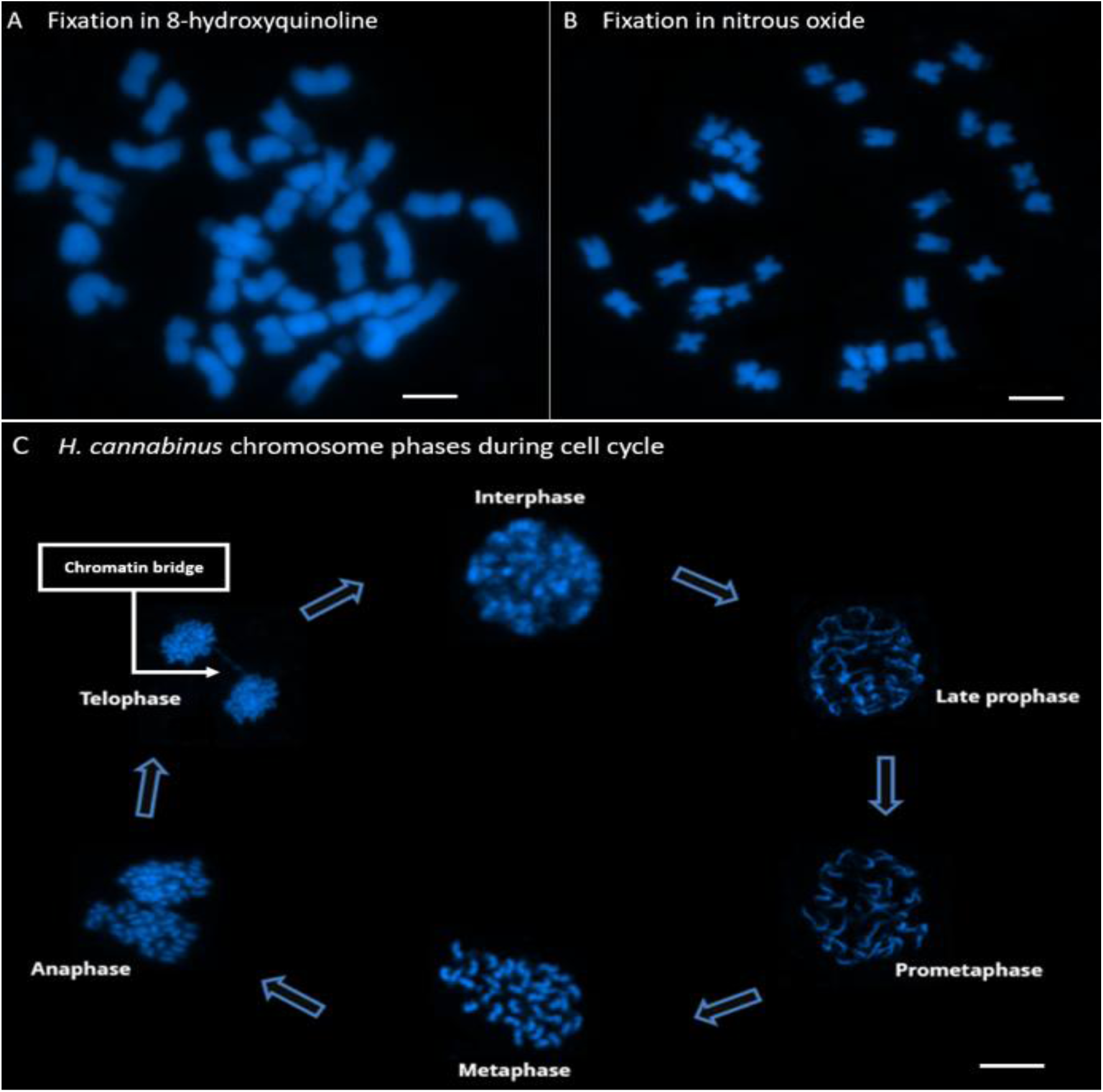
Morphology of kenaf (*H. cannabinus* EN-31) chromosomes stained with 4’,6-diamidino-2-phenylindole (DAPI) (A, B). Influence of chromosome preparation protocol on the morphology of kenaf chromosomes. (A) Fixation of root tips with 8-hydroxyquinoline leads to well-spread metaphases, usually without chromatid separation. (B) Fixation of root tips with nitrous oxide in a pressure chamber usually leads to slightly different chromosome condensation in metaphases and separation of chromatids. (C) Cell cycle phases of kenaf (*H. cannabinus)*. A chromatin bridge that formed during chromatid separation in the telophase is indicated with a white arrow. Scale bar - 5µm.

The chromosomes of the kenaf accessions in the current study were confirmed to be diploid having a configuration set of 2n=2x=36 each (Figure 1A, B; Table 1). All metaphase chromosomes of kenaf have a similar morphology with metacentric to sub-metacentric chromosomes. Nearly all chromosomes of kenaf showed visible, strong DAPI staining at the pericentromeric regions and weaker DAPI staining on the distal regions of many chromosome arms. In few instances (Figure 1A, B), weak DAPI staining was observed in distinct, intercalary regions (Figure 1A).

For the first time, kenaf chromosomes were followed along the mitotic cycle from interphase through to telophase (Figure 1C), showing the condensation and decondensation of chromatin. Kenaf metaphase chromosomes appeared differently in each phase of the cell cycle. Interestingly, in some nuclei, a chromatin bridge (Figure 1C, arrowed) was observed in the telophase stage of the mitotic cell cycle. This may be a possible indication of chromosomal instability in the genome of some kenaf breeding lines.

### Multi-color fluorescent *in situ* hybridization (FISH) reveals distinct rDNA loci and *Arabidopsis*-type telomeres

For species like kenaf with a paucity of chromosome markers, it was imperative to gain first insights with well-described rDNA probes. These were derived from the genes encoding the 5S (red fluorescence) and 35S (18S-5.8S-25S, green fluorescence) ribosomal RNAs as landmark probes for comparative cytogenetics (Figure 2A, B).

**Figure 2:**
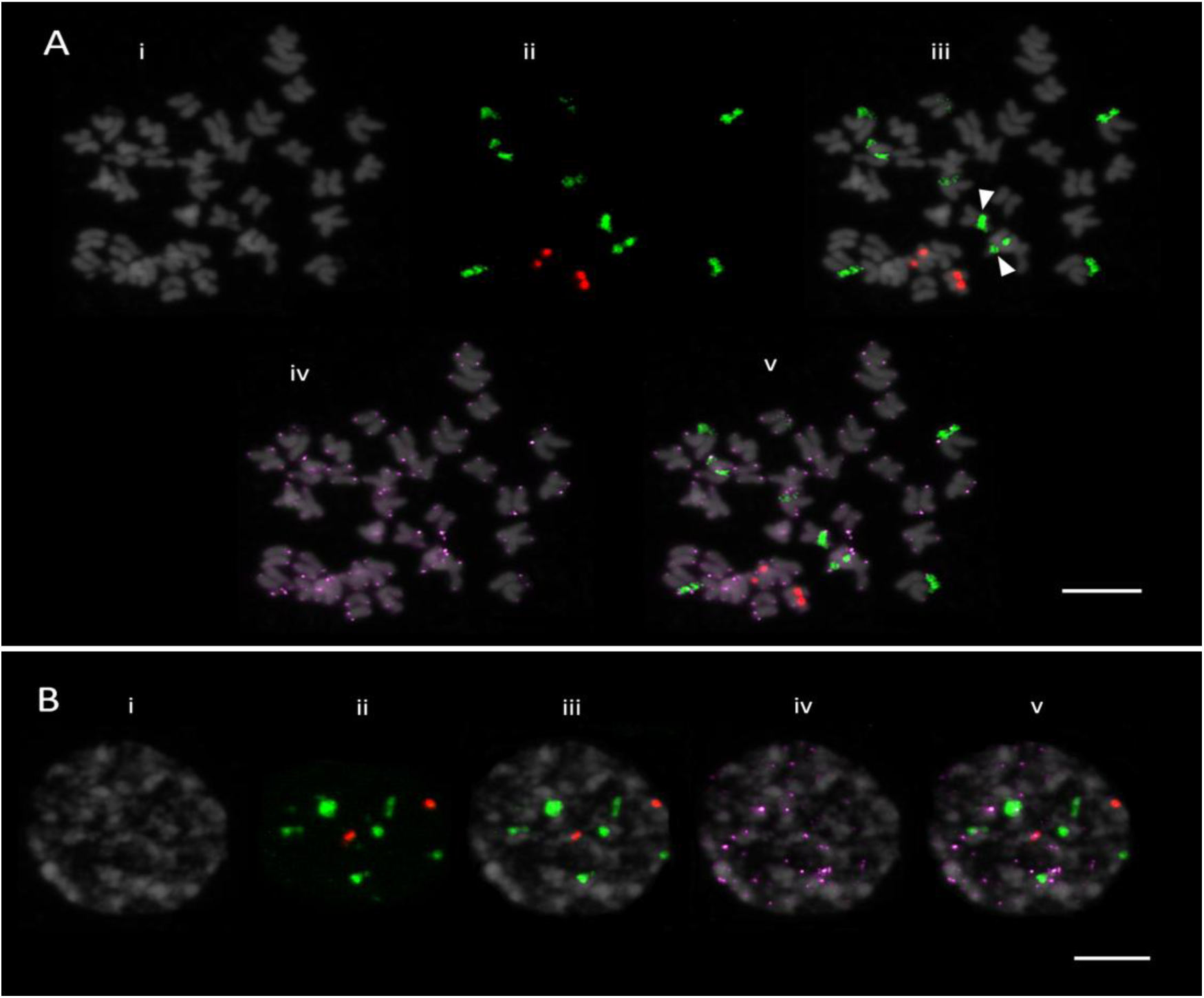
Fluorescent *in situ* hybridization of metaphase chromosomes (Figure 2A) and interphase nuclei (Figure 2B) using 18S rDNA, 5S rDNA and telomeric probes along chromosomes of kenaf (*H. cannabinus* EN-31). DAPI-stained DNA (grey) shows morphology of metaphase chromosomes and interphase nuclei (Figures 2A, B). Hybridization signals of 5S rDNA (red fluorescence) and 18S rDNA (green fluorescence) are indicated (Figure 2A-ii, iii; B-ii, iii). Arrowheads indicate interstitially localized 18S rDNA positions (Figure 2A iii). The telomeric probe pLT11 (purple fluorescence) localizes at terminal sites (Figure 2A-iv, B-iv). Scale bar - 5 µm.

FISH with rDNA probes showed hybridization signals of both 5S and 35S rDNA in the kenaf accessions analyzed. In all kenaf accessions, the 5S rRNA genes were localized in two sites (one pair of chromosomes) at the interstitial regions of the chromosomes (Figure 2A-iii, red fluorescence). For the 35S rRNA genes, most were located at six or eight sites (three pairs or four pairs of chromosomes), depending on the kenaf breeding line. Most 35S rDNA signals occur at the terminal ends of the kenaf chromosomes (Figure 2A-iii, green fluorescence), except for a signal pair in the interstitial regions (Figure 2A; arrow heads). The interstitial 35S signals occupy weakly DAPI-stained regions, indicating loosely packed euchromatin. Observed fluorescence of both 5S rDNA and 35S rDNA probes showed strong signals, but also a few weak signals of 35S (Figure 2A-ii).

To understand the nature of telomeres in the kenaf (*H. cannabinus*) accessions, FISH with the telomeric probe pLT11 binding to the tandem repeat “TTTAGGG” revealed the presence of *Arabidopsis*-type telomeric sequences at distal ends of all chromosome arms (Figure 2A iv, purple fluorescence). The fluorescence of the telomeric probe gave moderate and weak signals, especially compared to the strength of the 5S and 35S rDNA signals. Weak interstitial telomeric signals were also detected, potentially a sign of past chromosome rearrangements.

At higher resolution, 3-color FISH with 5S rDNA (red fluorescence), 35S rDNA (green fluorescence), and pLT11 (telomeric probe, purple fluorescence) to the interphase nuclei of the kenaf accessions allowed to investigate the compaction of the rDNAs (Figure 2B). Both rDNA types occupy the weakly DAPI-stained euchromatin. The 5S rDNA forms two loci (red), whereas the 35S rDNA loci form six major sites (green) in the interphase nuclei. Similarly, it was notable that the pLT11 probe showed both moderate and faint *Arabidopsis*-like telomeric fluorescence that was scattered across the kenaf interphase, without revealing any particular chromosome localization (Figure 2B).

### Comparative rDNA hybridization shows chromosomal variability in kenaf breeding lines

Summarizing the previous results, we have observed an informative rDNA pattern along kenaf chromosomes. Intriguingly, the 35S rDNA loci occur not only distally, but also in intercalary positions (Figure 2A-iii), likely resulting from recent chromosome rearrangements. Additionally, we have occasionally observed mitotic chromatin bridges in several ana- and telophases (Figure 1C; arrowed), potentially indicating chromosomal instability. To investigate, if the observations are a sign of chromosomal variability in kenaf, we then comparatively investigated nuclei of one landrace HC-21 and four breeding lines EN-31, TN-11, PN-51, HN-61 (Figure 3).

**Figure 3:**
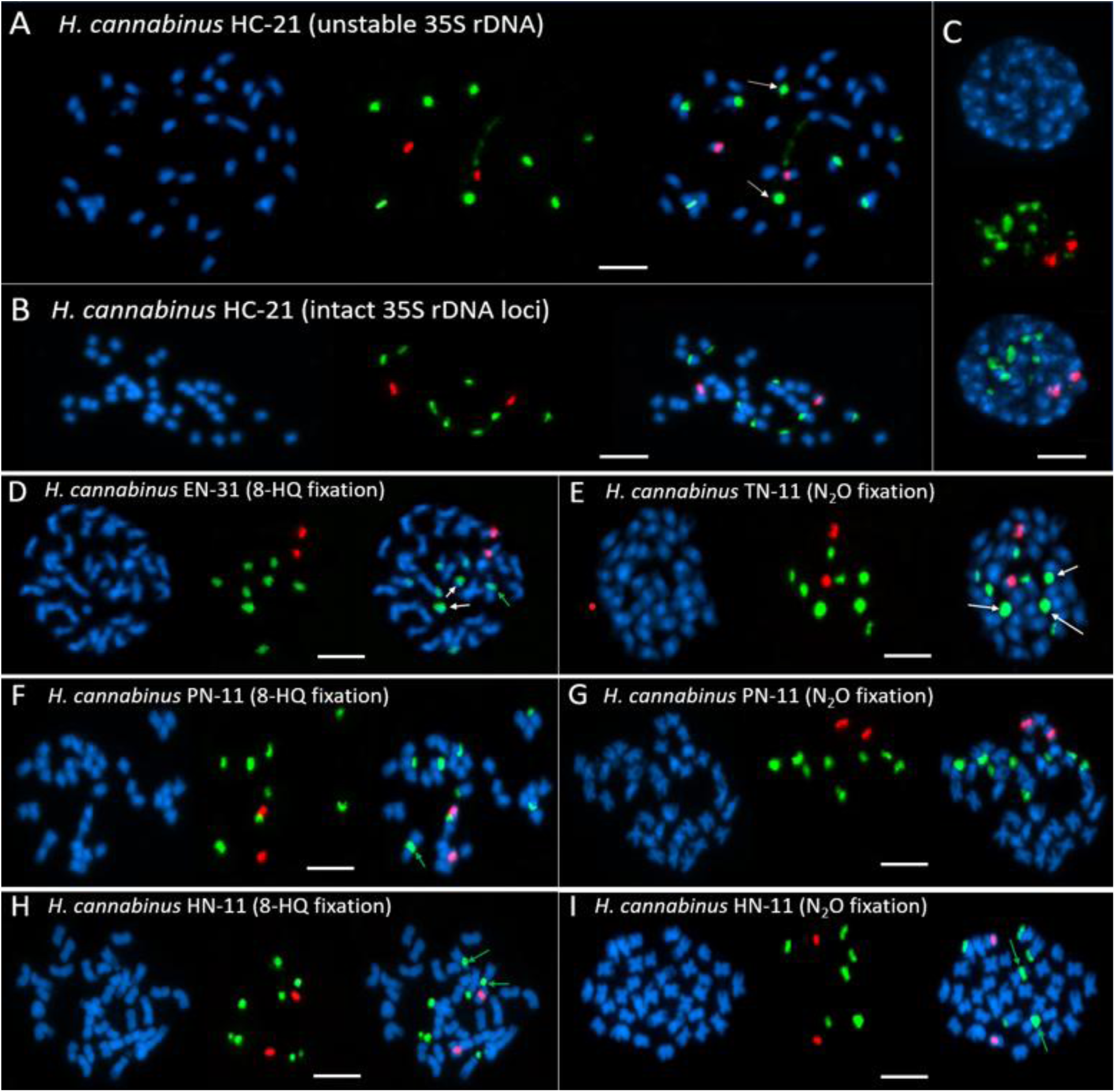
Comparison of the 5S and 35S rDNA loci number and position on chromosomes of kenaf accessions. Each panel shows the morphology of the kenaf chromosomes stained with DAPI (blue). Hybridization signals of 5S rDNA are visible as red fluorescence and the 35S rDNA probe hybridization indicates green fluorescence. Green arrows point out interstitially localized sites of 35S rDNA; white arrows indicate 35S rDNA signals on ripped-off nucleolar organizing regions (NORs). Scale bar - 5 µm. (A-C) Hybridization onto metaphase (A-B) and interphase (C) nuclei of the *H. cannabinus* line HC-21. (D-E) Hybridization onto *H. cannabinus* EN-31 (D) and TN-11 (E) chromosomes. (F-I) As the fixation protocol affects the chromosome morphology, we show 8-hydroxyquinoline-fixated (F, H) and nitrous oxide-fixated (G, I) chromosomes of the H. cannabinus breeding lines PN-11 (F-G) and HN-11 (H-I).

For the first time we established the application of FISH using the 5S rDNA and 35S rDNA probes to reveal the localization and the number of rDNA loci for five accessions of kenaf. The 5S rDNA (red fluorescence) is conserved in kenaf and consistently showed signals on two chromosomes across all the five kenaf accessions while the 35S rDNA (green fluorescence) is variable and unstable, showing signals on eight chromosomes or less frequently on six chromosomes of kenaf accessions.

The kenaf landrace HC-21 showed 5S rDNA signals in the interstitial positions of two chromosomes and also eight 35S rDNA signals at the sub-terminal and terminal positions of the chromosomes. The loose compaction of the 35S rDNA can lead to a ripping of this chromosomal area during chromosome preparation; this was often observed in HC-21 with the nucleolar organizing regions (NORs) ripped off from two chromosome arms (Figure 3A; white arrows), and in another panel, the complete metaphase showed the exact distribution of the rDNA regions along the kenaf chromosomes (Figure 3B). At higher resolution, the interphase nuclei of HC-21 showed both euchromatin (weak/dull) and heterochromatin (strong/bright) structures with DAPI staining showing six 35S rDNA and two 5S rDNA fluorescence signals (Figure 3C).

Kenaf breeding lines EN-31 and TN-11 having entire leaves and trilobed leaves, respectively, both showed similar fluorescent signals and intensities. The 5S rDNA localized interstitially on two chromosomes each for EN-31 and TN-11 (Figure 3D, E). In both EN-31 and TN-11 breeding lines, eight signals of the 35S rDNA fluorescence were observed, however, while five chromosomes of EN-31 were localized terminally/sub-terminally, one chromosome arm showed fluorescence at the interstitial position and two NORs were also ripped off (Figure 3D; green and white arrows). Similarly, for breeding line TN-11 also, five of the chromosome arms fluoresced with 35S rDNA and three NORs ripped off their chromosomes, indicating very open and hence more fragile chromatin at these regions (Figure 3E; white arrows).

For kenaf breeding lines PN-11 and HN-11 with penta-lobed leaves and hepta-lobed leaves respectively, two metaphases each were selected - one synchronized in 8-hydroxyquinoline (8-HQ) fixation and another with nitrous oxide (N_2_O) (Figure 3F-I). The metaphases of PN-11 in both 8-HQ and N_2_O treatments, had two chromosome arms each localized interstitially by the 5S rDNA (Figure 3F, G). Except for one 35S rDNA signal showing at interstitial site of one chromosome arm (green arrow), the other seven chromosomes were sub-terminally/terminally localized by 35S rDNA. The second panel PN-11, indicated a complete metaphase showing the exact distribution of the both 5S and 35S rDNA along the kenaf chromosomes (Figure 3G).

The breeding line HN-11 in both 8-HQ and N_2_O treatments, also had 5S rDNA localizing interstitially in two chromosome arms and three loci pairs of the 35S rDNA in the terminal and subterminal (Figure 3H, I; green arrows) sites of each of the differently treated metaphase chromosomes. Again, variability was observed in the signal pattern of the 35S rDNA; two chromosome arms of the kenaf HN-11 localized signals interstitially (red arrows), and the remaining signals were observed on four other chromosomes in both panels of HN-11 (Figure 3H, I).

## DISCUSSION

### Diploid kenaf has 36 small chromosomes with an informative rDNA pattern

High quality metaphase chromosome spreads are useful for chromosome number documentation and FISH experiment analysis (Andres & Kuraparthy 2013; Albert et al. 2023). The present study reports an optimized metaphase chromosomes preparation protocol that allows for reproducible high-quality kenaf metaphases using the maceration method combined with either 8-hydroxyquinoline (8-HQ) or nitrous oxide (N_2_O) pre-treatment and enzymatic digestion of root tips.

Pre-treatment using 8-HQ seems to be the most common fixation agent, however, 8-HQ fixation of root tips largely results in condensed chromosomes without separated chromatids and less frequently, condensed chromatids with poorly separated chromatids (Figure 1A). Nitrous oxide fixation on the other hand, results in properly condensed chromosomes with visibly separated sister chromatids (Figure 1B). Also, chromosomes pre-treated with N_2_O show discernible primary constrictions (centromere positions) compared to chromosomes pre-treated with 8-HQ which shows poorly visible centromere positions.

There is limited information on molecular cytogenetics of kenaf despite its economic value. In contrast, the techniques have been employed leading to the development of cytogenetic maps and karyotypes of major crops including soybean (Findley et al. 2010; 2017), potatoes (Dong et al. 2000), maize (Danilova & Birchler 2008; Mondin et al. 2014) and cotton (Gan et al. 2013; Lu et al. 2018; Liu et al. 2020).

Understanding chromosome data is the basis of cytogenetics (Braz et al. 2018; Han et al. 2015) and could be used to resolve inaccuracies that impair traditional classification (Jinyao et al. 2002; Zhu & Wang 2008). Accurate determination of chromosome number and ploidy of plant species plays an essential role in breeding and thus is efficient for hybridization schemes and promoting penetrance between species (Sakhanokho et al. 2020). Kenaf possesses small chromosome structures that are mostly metacentric to sub-metacentric with estimated lengths of 1.08-2.57 µm (Akpan & Hossain 1998). According to Silva et al. (2018), species with small chromosome structures present the challenge of uneasy discrimination of chromosomes, as well as difficulty in chromosome counts. The chromosome count for each kenaf accession in this study were 36 chromosomes conforming to the configuration of 2n=2x=36 (Zhang et al. 2020), implying the genotypes in Ghana were diploids. In addition, as a characteristic of many plant species, staining the chromosome of kenaf with DAPI indicated the pericentromeric regions of kenaf chromosomes were highly heterochromatic, whereas the distal regions of the chromosome arms contained more euchromatic DNA (Kim et al. 2005).

We report here the first account of chromosomal landmarks in kenaf, which we visualize using multicolor FISH. For these, we use the ribosomal genes as probes, which were complemented with a probe for the telomeres. Genes for the 5S rDNA and 35S rDNA are highly conserved in plant species and hence used as heterogeneous probes for physical mapping in chromosomes (Han et al. 2008). For many plant species, the 35S rDNA are more abundant, variable in number, unstable in position, thus they are sub-terminal/terminally located in chromosomes and less frequently found in interstitial sites and also ripped off from chromosome arms (Pedrosa et al. 2006; Roa & Guerra 2015; Garcia et al. 2017). Within the kenaf genome, variation in the number and position of 35S rDNA was observed among the accessions, with four of the accessions (HC-21, EN-31, TN-11, PN-11) bearing eight hybridized signals and one accession (HN-11) having six hybridized signals. The observed variation in the 35S rDNA within the same species rarely occurs, however, it has been previously reported in some plant species – *Hibiscus* species (Deepo et al. 2022), rice (Shishido et al. 2000), wild oats (Hayasaki et al. 2001), and *Aegilops* species (Raskina et al. 2004).

Similarly, the 5S rDNA is characteristically located in the interstitial sites of chromosome arms in many plant species (Roa & Guerra 2015; Garcia et al. 2017). In kenaf, the 5S rDNAs are highly conserved and hybridized interstitially on two chromosome arms of each accession, as have been characteristically reported in other crops such as common beans (Pedrosa et al, 2003), sugar beet (Schmidt et al. 2021) and *Phaseolus* species (Moscone et al. 1999). Notably, the location of these rRNA genes may not be essential for their functionality since many other species occur with single locus in other chromosome locations (Garcia et al. 2017).

Kenaf possesses *Arabidopsis*-like telomeres and these telomeric signals detected at the terminal ends of the kenaf chromosomes confirmed the integrity of the chromosomes and thus, contributed to accurate counting of the chromosomes. *Arabidopsis*-like telomeres are common in plants and have been previously reported in plants such as *Citrus clementina* (Deng et al. 2019), *Pinus* spp. (Hizume et al. 2002), *Podocarpus* spp. (Murray et al. 2002), and *Zea mays* (Koo et al. 2016).

### Kenaf chromosomes show signs of variability and maybe even instability

Instability in the chromosome of plants is reported to arise from sporadic whole or partial chromosome rearrangements (De Storme & Mason 2014). In the case of kenaf, the observation of chromatin bridges in the mitotic nuclei of some accessions, a sign of chromosomal instability, is being reported for the first time.

The variation detected in kenaf in terms of 35S rDNA localization in intercalary positions other than the terminal sites of chromosomes of 35S as well as the weak interstitial telomeric signals suggests that the kenaf 35S rDNA loci underwent repositioning, plausibly arising from structural “intra-chromosomal rearrangement mechanism” (Pedrosa et al. 2006) and/or by “breakage-fusion-bridge” phenomenon. Ribosomal RNA genes may also change their positions without changes in the order of surrounding genes (Dubcovsky & Dvorak 1995; Shishido et al. 2000) through transposon-mediated events (Raskina et al. 2004). These mechanisms are not rare as it has been studied in *Lotus japonicus* (Hayashi et al. 2001) and *Phaseolus vulgaris* (Pedrosa et al. 2006).

Again, the observed pattern of variability in the number of 35S rDNA across the kenaf accessions could be explained by a mechanism described by Dubcovsky & Dvorak (1995) as “dispersion-amplification-deletion” of rDNA repeats. Another plausible mechanism to explain the variability in the number of 35s rDNA loci in kenaf could be explained by ectopic (interlocus) recombination (Wendel et al. 1995). For species with distal 35S rDNA loci, unequal crossing over events occur and thus likely play major roles in loci size and number (Hanson et al. 1996). Results from other studies; *Paeonia* “garden peony” (Zhang & Sang 1999), *Aegilops speltoides* “goatgrass” (Raskina et al. 2004), and *Phaseolus vulgaris* “common beans” (Pedrosa et al. 2006) have supported the role of ectopic recombination, however, in the case of kenaf, extensive molecular cytogenetic studies must still be carried out on both cultivated and wild species to confirm the variation within 35S rRNA genes.

To better understand chromosomal, genomic and genetic variability in *Hibiscus* species, the insights gathered in kenaf could serve as reference for further investigations.

## CONCLUSION

We establish the cytogenetic analysis of kenaf (*H. cannabinus*) that allow to determine the number and structure of its chromosomes. With those, we report the chromosome number and rDNA distribution patterns of five Ghana kenaf accessions. Both 5S and 35S rDNAs are considered as strong cytogenetic markers, giving rise to eight to ten chromosomal landmarks, depending on the kenaf accession. Curiously, we observed several signs of chromosomal instability, such as the presence of chromatin bridges in the ana- and telophase as well as non-terminal and variable rDNA sites. Our study provides insights into the chromosomal organization of kenaf’s genome and thus provides a basis for understanding genomic variation in the kenaf germplasm and breeding lines.

## ACKNOWLEDGEMENTS

We thank Bayer Foundation for the Jeff-Schell Fellowship that provided research travel grants that supported this study. We are grateful to the KNUST Bast Fiber Research Group, Ghana for providing seeds for the kenaf accessions used for the study.

We also express our gratitude to all members of the Plant Cell and Molecular Biology laboratory at Technische Universität Dresden, for their contribution and invaluable insights with various aspects of this study.

## AUTHOR CONTRIBUTIONS

**Conceptualization:** NAA, AT, TH,

**Investigation:** NAA, AE, TH

**Methodology:** NAA, AE, SB, TH

**Resources:** AT, TH

**Writing (original draft):** NAA, TH

**Writing (review and editing):** NAA, AE, SB, AT, TH

